# Dopamine release in striatal striosome compartments in response to rewards and aversive outcomes during classical conditioning in mice

**DOI:** 10.1101/2024.09.18.613611

**Authors:** Tomohiko Yoshizawa, Makoto Funahashi

## Abstract

The striatum consists of two anatomically and neurochemically distinct compartments, striosomes and the matrix, which receive dopaminergic inputs from the midbrain and exhibit distinct dopamine release dynamics in acute brain slices. Striosomes comprise approximately 15% of the striatum by volume and are distributed mosaically. Therefore, it is difficult to selectively record dopamine dynamics in striosomes using traditional neurochemical measurements in behaving animals, and it is unclear whether distinct dynamics play a role in associative learning. In this study, we used transgenic mice selectively expressing Cre in striosomal neurons, combined with a fiber photometry technique, to selectively record dopamine release in striosomes during classical conditioning. Water-restricted mice could distinguish the conditioned stimulus (CS) associated with saccharin water from the air-puff-associated CS. The air-puff-associated CS evoked phasic dopamine release only in striosomes. Furthermore, air puff presentation induced dopamine release to striosomal neurons but suppressed release to putative matrix neurons. These findings suggest that dopamine is released in a differential manner in striosomes and the matrix in behaving animals and that dopamine release in striosomes is preferentially induced by the air-puff-associated CS and air puff presentation. These findings support the hypothesis that striosomal neurons play a dominant role in aversive stimuli prediction.

**Highlights:**

- Dopamine (DA) release was successfully measured in striosomes of behaving mice.
- A conditioned stimulus for aversive outcomes induced DA release in striosomes.
- In striosomes, rewarding and aversive unconditioned stimuli triggered DA release.

## Introduction

Humans and animals can learn new behaviors in unfamiliar environments by exploring and memorizing sensory cues or actions that lead to positive or negative outcomes. The striatum, a major cortical input site in the basal ganglia, plays an important role in predicting outcomes. Many lines of research, including functional brain imaging (O’Doherty et al., 2004; Tanaka et al., 2004) and electrophysiological recordings (Ito and Doya, 2009, 2015; Kim et al., 2009; Samejima et al., 2005), have shown that neural activity in the striatum reflects state and action values, which are predictions of forthcoming rewards from the current sensory state and possible action, respectively. The striatum is composed of a patchwork of two types of compartments, striosomes (also known as paches) and the matrix. Striosomes comprise only 15% of the striatum and are scattered mosaically. Striosomal neurons have inputs from the limbic cortex, and GABAergic neurons in striosomes have been revealed to have monosynaptic projections to dopaminergic neurons in the substantia nigra pars compacta (SNc) (Gerfen, 1984). The spike activity of dopaminergic neurons in the SNc encodes reward prediction error, which is defined as the discrepancy between the actual and predicted rewards (Schultz et al., 1997). Therefore, it is hypothesized that striosomes contribute to reward prediction (Barto, 1995; Doya, 2000). This hypothesis was supported by striosome-selective *in vivo* calcium imaging obtained while mice performed classical and operant conditioning tasks (Bloem et al., 2017; Yoshizawa et al., 2018). We found that striosomal neurons showed reward- or air puff-predictive activities; therefore, they encoded the value of the conditioned stimulus (CS). A principal distinction between striosome and non-selective imaging, including the putative matrix, was the greater proportion of air-puff-predictive neurons observed in striosomes than in the putative matrix. Neurons in the dorsal striatum receive glutamatergic and dopaminergic inputs from the cortex and the SNc, respectively. Dopamine (DA) modulates cortico-striatal synaptic plasticity (Reynolds et al., 2001), and its release into the striatum plays a critical role in classical conditioning (Iino et al., 2020; Yagishita et al., 2014). Some dopaminergic neurons have been shown to project preferentially to the striosomes and matrix compartments (Matsuda et al., 2009). Our previous study demonstrated significant roles for striosomal neurons in air puff prediction (Yoshizawa et al., 2018). In combination, striatal neurons in the striosomes and matrix compartments may receive distinct reinforcement signals through compartment-specific DA dynamics during classical conditioning. Although some studies have reported different DA dynamics between striosomes and the matrix in brain slices (Krebs et al., 1994; Krebs et al., 1991), it is unclear whether this difference is observed in behaving animals. The present study used a fiber photometry technique to examine DA dynamics in mouse striosomes during classical conditioning.

## Materials and Methods

### Subjects

The Hokkaido University Animal Use Committee approved this study. Male Pdyn-IRES-Cre mice (129S-Pdyn(tm1.1(Cre)/Mjkr)/LowlJ, The Jackson Laboratory Cat# 027958) were housed individually under a 12/12 h light/dark cycle (lights on at 7 A.M.; off at 7 P.M.). Experiments were performed during the light phase. Water was restricted to 1–2 mL/day for 2 days before experimental initiation and during the experimental period. Food was provided *ad libitum* for the entire period.

### Behavioral task

The mouse head and body were restricted using a head plate and a metal tube, respectively (Fig. 1A). From 3–5 days before the DA recording sessions, mice performed pretraining sessions in which an 8 kHz tone was presented for 2 s, followed by presentation of a drop of 0.1% saccharin water (4 µL). In the DA recording sessions, mice were presented with one of two tones (Tone A or Tone B) for 2 s, followed by an outcome in each trial (Fig. 1B). Tone A and Tone B were associated with different outcomes: a drop of 0.1% saccharin water (4 µL) and an air puff delivered to the animal’s face, respectively. An air puff is generally used as an aversive stimulus in rodents and is known to cause avoidance behaviors such as predictive eye blinks (Cohen et al., 2012; Heiney et al., 2014). The tones were presented randomly but with 80% probability for Tone A and 20% probability for Tone B. In six mice, Tone A and Tone B were 6 and 10 kHz pure tones, respectively. In the other four mice, the tone and outcome combination was reversed to eliminate a possibility that mice innately preferred a specific tone frequency. Inter-trial intervals ranging from 10 to 16 s were pseudorandomly selected. A daily session consisted of 200 trials (saccharin water: 160 trials, air puff: 40 trials). Licks were detected by interruptions of an infrared beam placed in front of the water tube.

**Figure 1.**
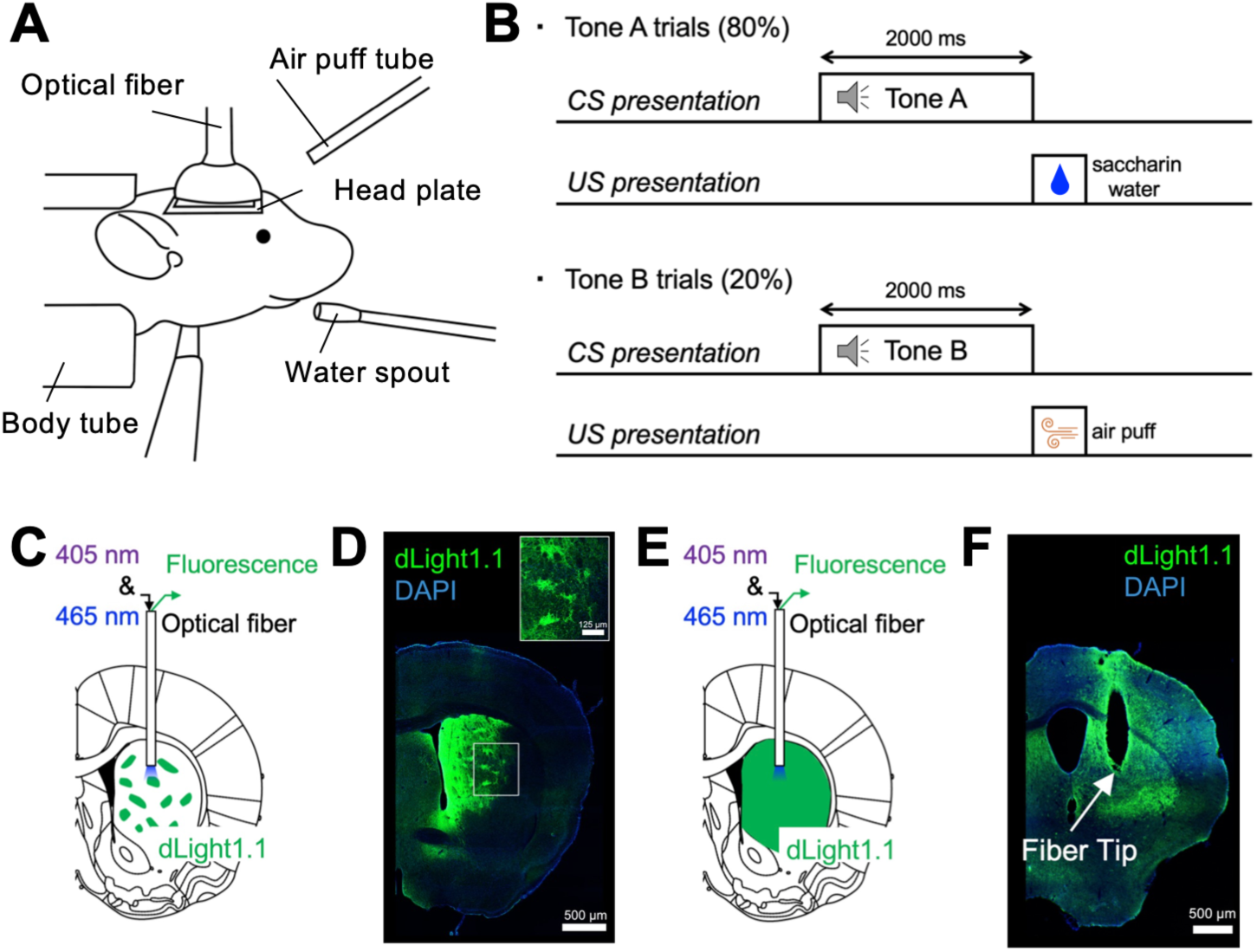
Dopamine measurement in striosomes during classical conditioning. **A.** Schematic illustration of the behavioral apparatus. The mouse head and body were restricted by a metal frame and tube. The waterspout and air puff tube were set in front of the mouth and eyes. Spout-licking behaviors were monitored using an infrared sensor. An optical fiber was connected to the optical probe implanted in the dorsomedial striatum (DMS) for fiber photometry of dLigh1.1. **B.** Time course of a classical conditioning task. Each trial began with a 2-s presentation of one of two tones, followed by an outcome. Tone A and B were associated with a drop of saccharin water and an air puff, respectively. **C.** Schematic illustration of the measurement of dopamine (DA) dynamics in striosomes. dLight1.1 was selectively expressed in striosomes by injection of AAV2/9-EF1α-DIO-dLight1.1 into the striatum of prodynorphin-Cre mice. An optical fiber was implanted in the DMS to measure the DA-dependent fluorescence of dLight1.1 excited by a 465 nm LED using the fiber photometry technique. **D.** Histological images of Cre-dependent dLight1.1-expressing neurons in the striatum. dLight1.1 was mosaically expressed in the striatum. **E.** Schematic illustration of measurement of DA dynamics from the putative matrix. dLight1.1 was expressed in both the matrix and striosomes by injection of AAV2/9-CAG-dLight1.1 into the striatum of prodynorphin-Cre mice. Because the majority of striatal neurons belong to the matrix, the dLight1.1 signal was predominantly derived from the matrix. **F.** Histological images of dLight1.1-expressing neurons within the striatum and optical fiber tract.

### Surgery

Mice were anesthetized with isoflurane (1.0%–4.0%) and placed in a stereotaxic frame. The skull was exposed, and a hole was drilled in the skull. For fiber photometry recording, AAV2/9-EF1α-DIO-dLight1.1 (left hemisphere: two mice, right hemisphere: three mice, PT1139, BrainVTA, Wuhan, China, Fig. 1C, D) or AAV2/5-CAG-dLight1.1 (left hemisphere: one mouse, right hemisphere: four mice, 111067-AAV5, Addgene, Watertown, MA, USA, Fig. 1E, F) was injected into the dorsomedial striatum (DMS) (anteroposterior, AP: +0.5, mediolateral, ML: 1.75, dorsoventral, DV: 2.85 mm from the brain surface, volume: 400 nL) using a microsyringe pump (Legato100, Kd Scientific, Holliston, MA, USA). After AAV injection, an optical fiber (diameter: 400 μm, length: 5 mm, MFC_400/430-0.66_5mm_ZF1.25(G)_FLT, Doric, Quebec, Canada) was implanted 200 μm above the AAV injection coordinates (AP: +0.5, ML: 1.75, DV: 2.65 mm from the brain surface). The optical fiber was fixed with adhesive dental cement (Super bond, Sun Medical, Shiga, Japan). A head plate (CF-10, Narishige, Tokyo, Japan) was fixed with pink dental cement (Unifast 2, GC, Tokyo, Japan). The dental cement has confirmed effects on brain tissue (Yoshizawa and Funahashi, 2020). Analgesics and antibiotics were applied postoperatively as required (meloxicam, 1 mg/kg s.c.; 0.1% gentamicin ointment, *ad usum externum*).

### Fiber photometry

We referred to a previously published fiber photometry recording protocol for mice (Patel et al., 2020). The fiber photometry system consisted of two excitation channels. A 465 nm LED (CLED_465, Doric) and a 405 nm LED (CLED_405, Doric) were driven by an LED controller (LEDD_4, Doric) to obtain a DA-dependent signal and a DA-independent isosbestic signal, respectively. These LEDs were alternately turned on and off at 13.3 Hz. Fluorescence from dLight1.1 and isosbestic fluorescence were directed through dichroic mirrors (iFMC6_IE(400-410)_E1(460-490)_F1(500-540)_E2(555-570)_F2*(580-680)_S, Doric) and were acquired using a photodetector. The signals were passed through a 10× amplifier and were sampled at 1 kHz with a data acquisition system (Power1401, Cambridge Electronic Design, Cambridge, UK). The acquired photometry signals were processed using custom-written MATLAB code (MATLAB R2018a, Mathworks, Natick, MA, USA). First, the signals were downsampled to 13.3 Hz for further analysis. A fitting curve was estimated and subtracted from the original signal to remove exponential and linear signal decay during the recording session. A linear fit was applied to align the 405-nm signal to the 465-nm signal, and then the fitted 405-nm signal was subtracted from the 465-nm channel and divided by the fitted 405-nm signal to calculate ΔF/F values. The ΔF/F time-series trace was normalized using z-scores to account for data variability across animals and sessions.

### Immunohistochemistry

After all experiments were completed, mice were deeply anesthetized with pentobarbital sodium and then perfused with 4% paraformaldehyde. The brains were carefully removed so that the optical fibers would not cause tissue damage, post-fixed in 4% paraformaldehyde at 4 °C overnight, and then transferred to a 30% sucrose/phosphate-buffered saline (PBS) solution at 4 °C until they sank to the bottom. Coronal sections including the striatum were cut at a thickness of 50 μm on a freezing microtome (REM-710; Yamato, Saitama, Japan). Free-floating sections were washed in PBS for 15 min and placed in blocking buffer containing 10% normal donkey serum (017-000-121, Jackson ImmunoReserch Laboratories, West Grove, PA, USA) and 0.1% Triton X-100 in PBS for 1 h at room temperature. Sections were incubated in chicken anti-GFP primary antibody (GFP-1010, Aves Labs, Davis, CA, USA) diluted 1:500 in blocking buffer overnight at 4 °C. Afterward, the sections were washed four times for 15 min in PBS and temporarily stored at 4 °C. The sections were then incubated in donkey anti-chicken Alexa Fluor 488 secondary antibody (703-545-155, Jackson ImmunoResearch Laboratories) diluted 1:500 in blocking buffer overnight at 4 °C. The next morning, the sections were washed four times for 15 min in PBS, mounted on glass slides, and coverslipped with VECTASHIELD Mounting Medium with DAPI (Vector Laboratories, Newark, CA, USA). A fluorescence microscope (Eclipse Ci-L, Nikon, Tokyo, Japan) was used to inspect stained tissue, and images were obtained using NIS-Elements software (NIS-Elements D, Nikon).

### Experimental design and statistical analysis

The analyses included 6000 behavioral and neural trials recorded over a total of 30 sessions in 10 mice. Appropriate statistical tests were used when applicable, i.e., paired or unpaired *t*-tests and Pearson correlation analysis with or without Bonferroni’s multiple comparisons test. Differences were considered statistically significant when *p* < 0.05.

## Results

### Tone discrimination through classical conditioning

Head-fixed mice were classically conditioned with two sound cues associated with an unconditioned stimulus (US), including either saccharin water or an air puff, with the same apparatus (Fig. 1A) used in our previous study (Yoshizawa et al., 2018). As shown in Fig. 1B, in each trial, Tone A or Tone B was presented for 2 s as a CS, followed by a drop of 0.1% saccharin water or an air puff. To assess whether mice predicted forthcoming saccharin water, we counted the number of licks toward the waterspout. In Tone A trials, mice began to lick during the tone presentation period prior to water presentation. In Tone B trials, however, they did not lick during this period (Fig. 2A). The number of licks during the CS period was significantly greater in Tone A trials than in Tone B trials from day 1 to day 3 of conditioning (day 1: 3.5 ± 0.66, Tone A, and 1.1 ± 0.42, Tone B, *p* = 0.019; day 2: 4.7 ± 0.77, Tone A, and 1.4 ± 0.41, Tone B, *p* = 0.0019; day 3: 4.8 ± 0.93, Tone A, and 2.3 ± 0.58, Tone B, *p* = 0.015, mean ± SEM, paired *t*-test, Fig. 2B). These results indicate that mice can distinguish Tone A from Tone B and predict forthcoming water from Tone A by associative learning. These results are consistent with our previous study on odor classical conditioning (Yoshizawa et al., 2018).

**Figure 2.**
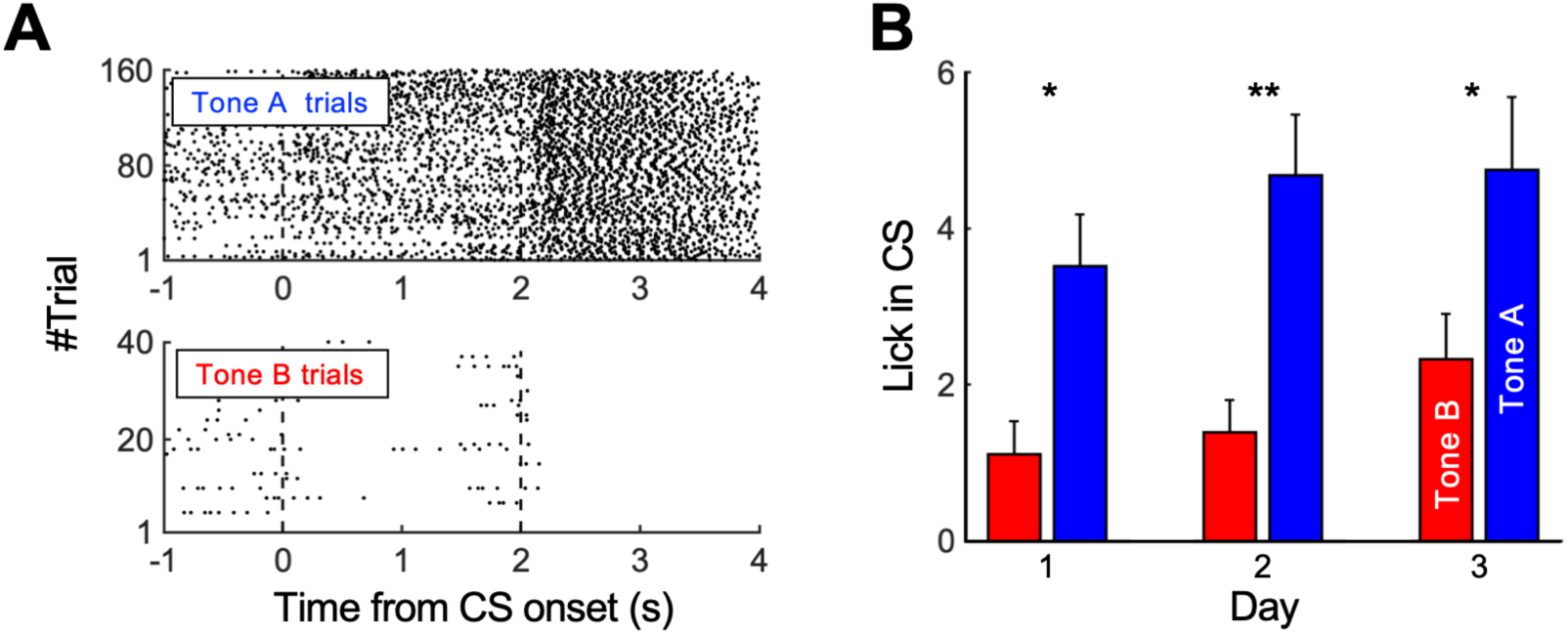
Mice distinguished Tone A from Tone B. **A.** An example of spout-licking behaviors on day 3. In Tone A trials, spout-licking behaviors were observed during the presentation of Tone A. However, in Tone B trials, these behaviors were not observed during the presentation of Tone B. Black dots indicate the timing of spout-licking behaviors. **B.** Average number of spout-licking behaviors during conditioned stimulus (CS) periods in all 10 mice. Error bars indicate the SEM. **: *p* < 0.01. *: *p* < 0.05, paired *t*-test.

### Dopamine dynamics in striosomes during classical conditioning

Using the fiber photometry technique, we evaluated the DA dynamics of striosomes during 3 days of behavioral sessions (Fig. 1C). Transgenic mice (Pdyn-IRES-Cre) selectively expressing Cre in their striosomal neurons were employed (Evans et al., 2020; Xiao et al., 2020). To selectively express the genetic DA sensor dLight1.1 (Patriarchi et al., 2018) in striosomal neurons, AAV2/9-EF1α-DIO-dLight1.1 was injected into the DMS (Fig. 1D).

In the Tone A trials, the first small DA signal elevations (z-scored dLight1.1 fluorescence) were observed at the onset of Tone A (representative mouse, Fig. 3A, B). The second large DA signal elevations were observed when the mouse drank saccharin water. Although the correlation between the trial order and the DA response to Tone A was not significant over the 3 days of behavioral sessions (r_A_ = −0.031, *p* = 0.49, Fig. 3C), the correlation between the trial order and DA response to saccharin water was significantly positive (r_water_ = 0.33, *p* = 1.0e−13, Fig. 3D), indicating a learning effect on DA release in response to saccharin water intake. In Tone B trials, similar to Tone A trials, the first and second elevations in the DA signal were observed at the onset of Tone B and the receipt of air puffs, respectively (Fig. 3E). The second DA release gradually diminished over 3 days (Fig. 3F). Although the correlation between the trial order and DA response to Tone B was not significant over the 3 days of behavioral sessions (r_B_ = −0.089, *p* = 0.33, Fig. 3G), the correlation between the trial order and DA response to air puffs was significantly negative (r_air_ = −0.49, *p* = 1.6e−08, Fig. 3H), indicating a learning effect on DA release in response to air puff presentation.

**Figure 3.**
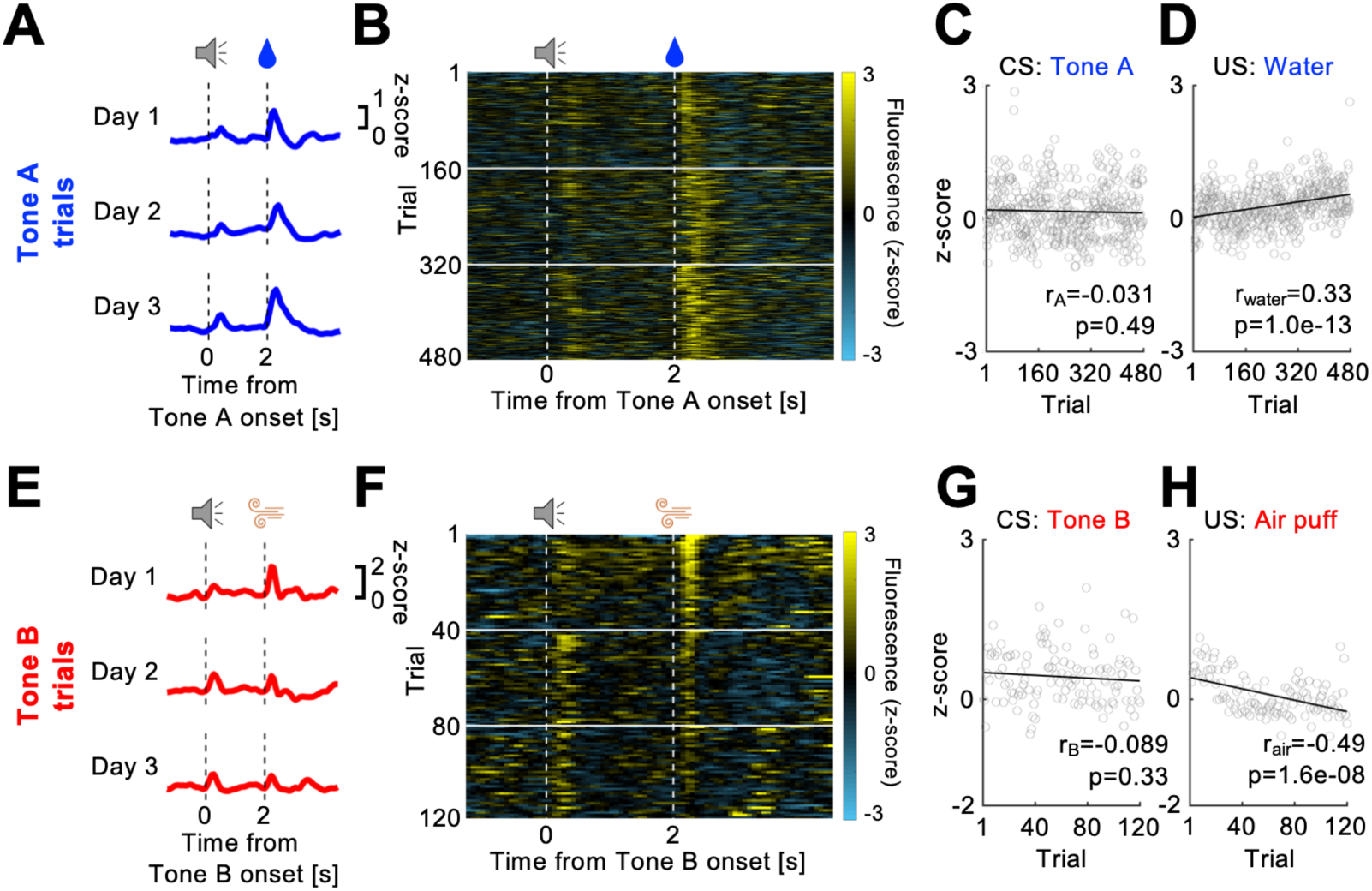
Dopamine dynamics in striosomes during classical conditioning. **A.** Representative DA dynamics in striosomes during Tone A trials. Each blue line indicates the average fluorescence in Tone A trials of different days. Saccharin water intake induced DA release over 3 days. **B.** The fluorescence in each Tone A trial is presented as a heat map. **C.** Correlation between the number of Tone A trials and DA release during Tone A presentation. The average fluorescence during the 1 s from Tone A onset is plotted against the number of Tone A trials. Gray circles and the black line indicate the average fluorescence in each trial and the regression line, respectively. Pearson correlation analysis. **D.** Correlation between the number of Tone A trials and DA release during the 1.5 s after saccharin water intake. DA release in response to saccharin water was positively correlated with experience of Tone A trials. **E.** Representative DA dynamics in striosomes during Tone B trials. Each red line indicates the average fluorescence in Tone B trials on different days. Both Tone A and air puffs induced DA release in striosomes. **F.** The fluorescence in each Tone B trial is presented as a heat map. **G.** Correlation between the number of Tone B trials and DA release during Tone B presentation. The average fluorescence during the 1 s after Tone B onset is plotted against the number of Tone B trials. **H.** Correlation between the number of Tone B trials and DA release during the 1.5 s after air puff onset. DA release in response to the air puff showed a significant negative correlation with experience of Tone B trials.

### Dopamine dynamics in the putative matrix during classical conditioning

We next examined how DA was released in the matrix during classical conditioning. We prepared mice expressing dLight1.1 in both striosomes and the matrix by injecting AAV2/5-CAG-dLight1.1 into the DMS (Fig. 1E, F). Because approximately 85% of the striatum consists of matrix, the dLight1.1 signal was predominantly derived from the matrix. Fig. 4A shows a representative example of a DA signal activated by saccharin water intake. Saccharin water consistently induced DA release on all 3 days (Fig. 4B). The correlations between the trial order and the DA response to Tone A or saccharin water intake were not significant over the 3 days of behavioral sessions (r_A_ = 6.5e−03, *p* = 0.89; r_water_ = −0.020, *p* = 0.66, Fig. 4C, D). In Tone B trials, presentation of air puffs induced DA release on day 1 (Fig. 4E). This DA release became weaker on day 2 and then eventually dropped below zero on day 3 (Fig. 4F). Although the correlation between the trial order and DA response to Tone B was not significant (r_B_ = −0.11, *p* = 0.25, Fig. 4G), the correlation between the trial order and DA response to air puffs was significantly negative (r_air_ = −0.20, p = 0.032, Fig. 4H), indicating a learning effect on DA release in response to air puffs.

**Figure 4.**
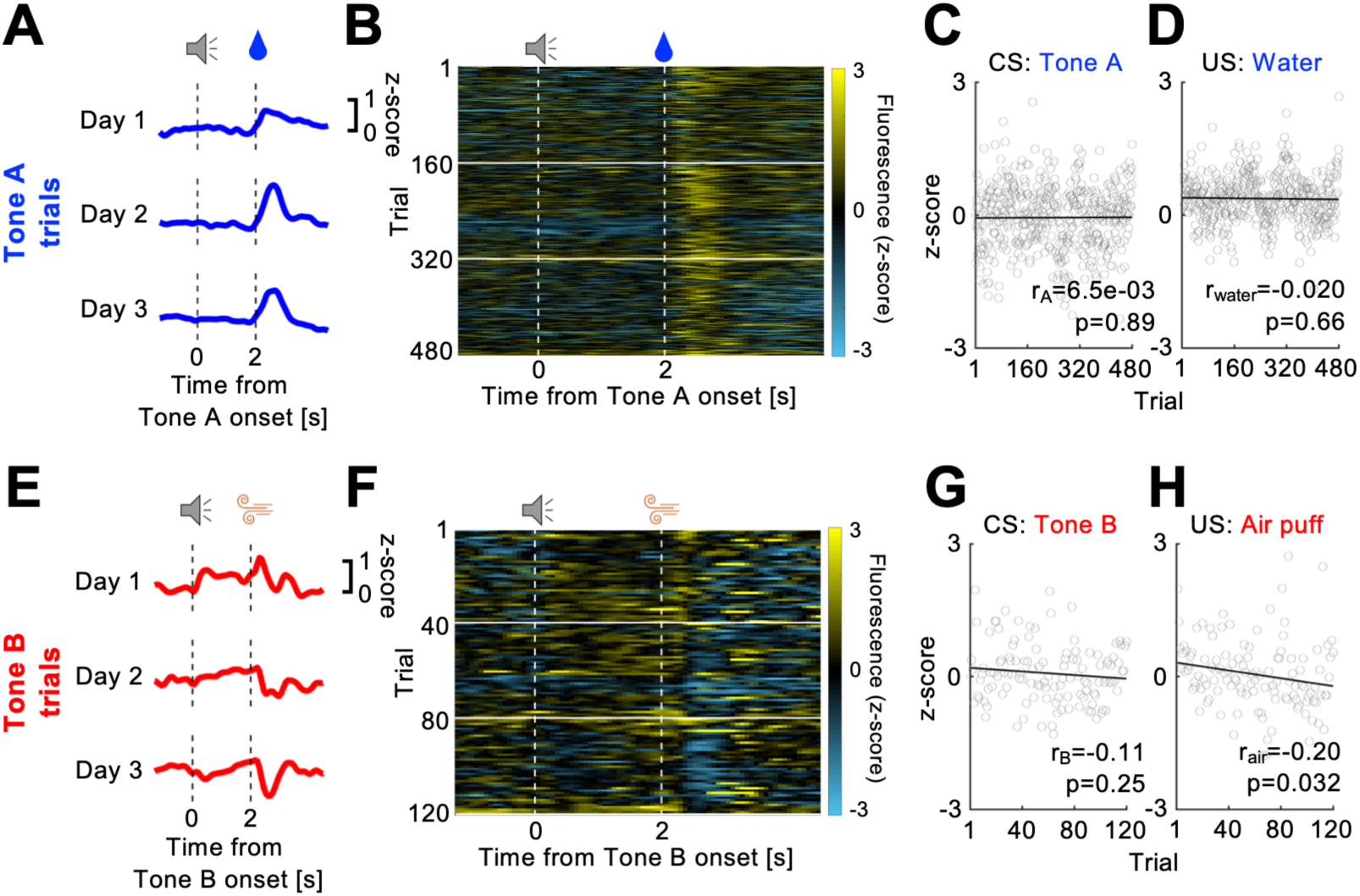
DA release in the putative matrix during classical conditioning. **A.** Representative DA dynamics in the putative matrix during Tone A trials. Each blue line indicates the average fluorescence in Tone A trials on different days. Saccharin water intake induced DA release over 3 days. **B.** The fluorescence in each Tone A trial is presented as a heat map. **C.** Correlation between the number of Tone A trials and DA release during Tone A presentation. The average fluorescence during the 1 s after Tone A onset is plotted against the number of Tone A trials. Gray circles and the black line indicate the average fluorescence in each trial and the regression line, respectively. Pearson correlation analysis. **D.** Correlation between the number of Tone A trials and DA release during the 1.5 s after saccharin water intake. **E.** Representative DA dynamics in the putative matrix during Tone B trials. Each red line indicates the average fluorescence in Tone B trials on different days. In contrast to striosomes, the air puff depressed DA release in the putative matrix. **F.** The fluorescence in each Tone B trial is presented as a heat map. **G.** Correlation between the number of Tone B trials and DA release during Tone B presentation. The average fluorescence during the 1 s after Tone B onset is plotted against the number of Tone B trials. **H.** Correlation between the number of Tone B trials and DA release during the 1.5 s after air puff onset. DA release in response to air puffs decreased significantly with increasing number of Tone B trials.

### Comparison of DA dynamics between striosomes and the putative matrix

To quantitatively examine differences in DA dynamics between striosomes and the putative matrix, we averaged the dLight1.1 fluorescence of all mice in each group. The DA release in response to Tone B (1 s duration from Tone B onset) was significantly larger than the baseline signal (averaged z-score during the 2 s before Tone B onset) over 3 days in striosomes, whereas this difference was not significant in the putative matrix (day 1, Fig. 5A, D: *p* = 0.023, striosomes, *p* = 0.38, putative matrix; day 2, Fig. 5B, E: *p* = 7.7e−06, striosome, *p* = 0.71, putative matrix; day 3, Fig. 5C, F: *p* = 1.6e−09, striosomes, *p* = 0.87, putative matrix, paired *t*-test, followed by Bonferroni correction, Fig. 5G). Moreover, in striosomes, Tone B responses were significantly larger than those for Tone A trials on days 2 and 3 (day 2: −0.022 ± 0.031, reward, and 0.075 ± 0.056, air puff, *p* = 0.044; day 3: 0.040 ± 0.037, reward, and 0.16 ± 0.061, air puff, *p* = 0.032, unpaired *t*-test, followed by Bonferroni correction).

**Figure 5.**
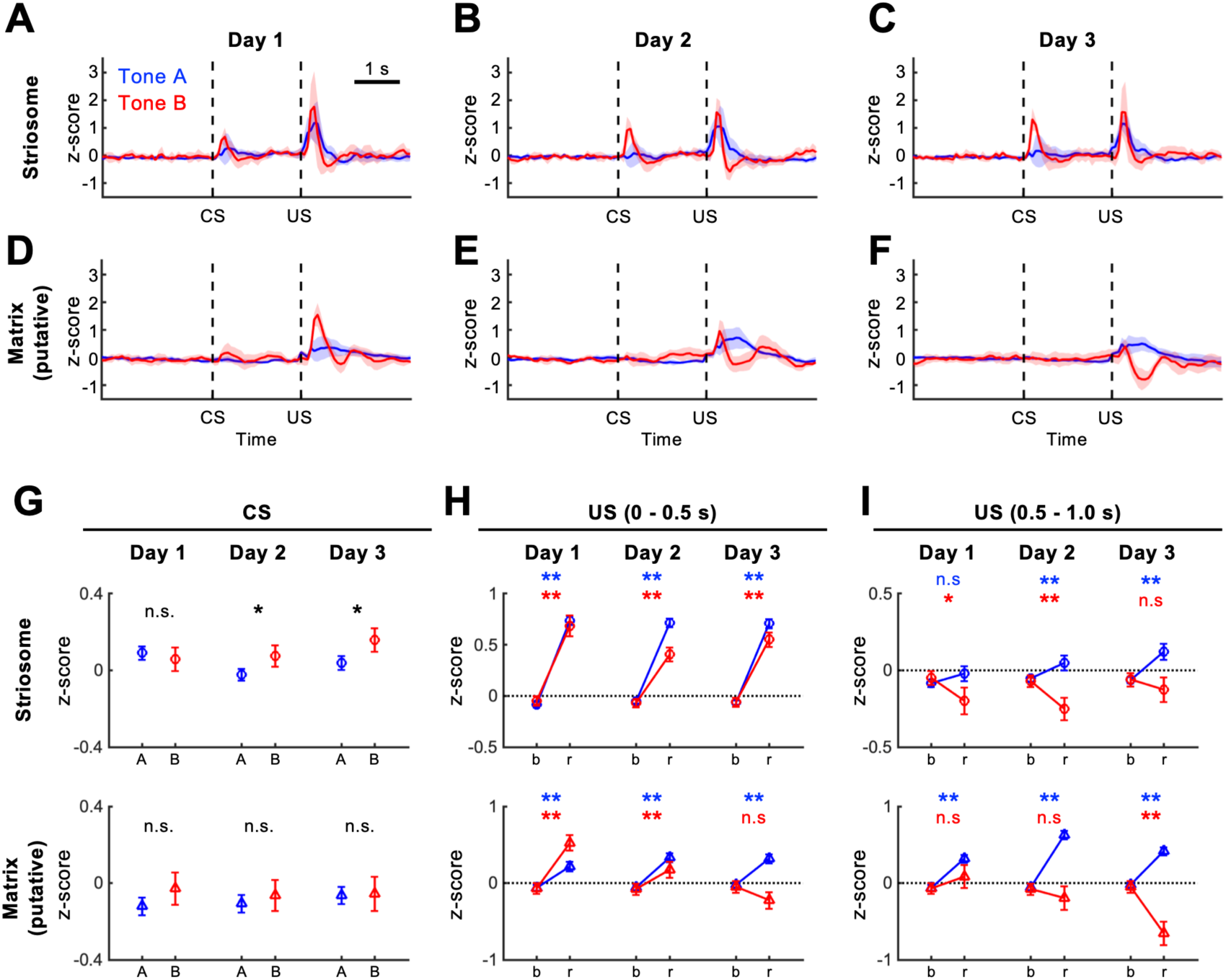
DA dynamics were significantly different between striosomes and the putative matrix. **A–C.** Population DA dynamics in striosomes (n = 5 mice) on day 1 (**A**), day 2 (**B**), and day 3 (**C**). In striosomes, DA was released in response to Tone B over 3 days. Both saccharin water and air puffs induced DA release in striosomes. Blue and red shadows indicate 95% confidence intervals. **D–F.** Population DA dynamics in the putative matrix (n = 5) on day 1 (**D**), day 2 (**E**), and 3 (**F**). In the putative matrix, saccharin water induced DA release, whereas the air puff gradually decreased DA release over 3 days. **G.** Quantitative analysis of DA release in response to the CS. A: Tone A, B: Tone B. Error bars indicate the SEM. *: *p* < 0.05, n.s.: *p* ≥ 0.05, unpaired *t*-test followed by Bonferroni correction. **H, I.** Quantitative analysis of DA release during the first (0–0.5 s after unconditioned stimulus (US) onset, H) and second half (0.5–1 s after US onset, I) of the US period. **: *p* < 0.01, *: *p* < 0.05, n.s.: *p* ≥ 0.05, paired *t*-test followed by Bonferroni correction. b: baseline (averaged z-score during 2 s before CS onset), r: response to US.

We next analyzed the DA responses to USs. In striosomes, both saccharin water and air puffs induced significantly more DA release than that released at baseline (day 1: *p* = 6.0e−150, reward, *p* = 1.3e−20, air puff, day 2: *p* = 1.1e−127, reward, *p* = 5.0e−21, air puff, day 3: *p* = 2.2e−124, reward, *p* = 3.0e−25, air puff, paired *t*-test, followed by Bonferroni correction) during the early US period (0 to 0.5 s after US onset), reflecting the salience of stimuli (Fig. 5H). In the putative matrix, DA release in response to air puffs appeared biphasic with a positive to negative response (Fig. 5D–F). The DA response in the early positive phase (0 to 0.5 s after air puff onset) gradually decreased each day (Fig. 5H). The DA response in the late negative phase (0.5 to 1.0 s after air puff onset) was significantly lower than the baseline signal on day 3 (*p* = 2.3e−11, Fig. 5I). The DA response to saccharin water was significantly larger than the baseline signal during the entire US period and on all 3 days (Fig. 5H, I). These results suggest that DA signals to striosomes more strongly reflect information regarding Tone B and air puffs than those to the matrix.

We finally compared the learning effects on DA release between striosomes and the putative matrix. Population analysis of the correlation coefficients in Tone A trials revealed that both r_A_ and r_rwd_ were not significantly different between striosomes and the putative matrix (r_A_: −0.076 ± 0.043, striosome, 0.028 ± 0.019, putative matrix, *p* = 0.058, Fig. 6A; r_water_: −0.053 ± 0.10, striosome, −0.012 ± 0.071, putative matrix, *p* = 0.75, Fig. 6B, mean ± SEM, unpaired *t*-test). In Tone B trials, the r_B_ coefficient of striosomes was larger than that of the putative matrix (striosome: 0.092 ± 0.055, putative matrix: −0.049 ± 0.023, *p* = 0.046, Fig. 6C), whereas the rair correlation coefficients did not differ (striosome: −0.13 ± 0.13, putative matrix: −0.34 ± 0.046, *p* = 0.16, Fig. 6D). These results indicate that Tone B learning has a stronger effect on DA dynamics in striosomes than on those in the putative matrix.

**Figure 6.**
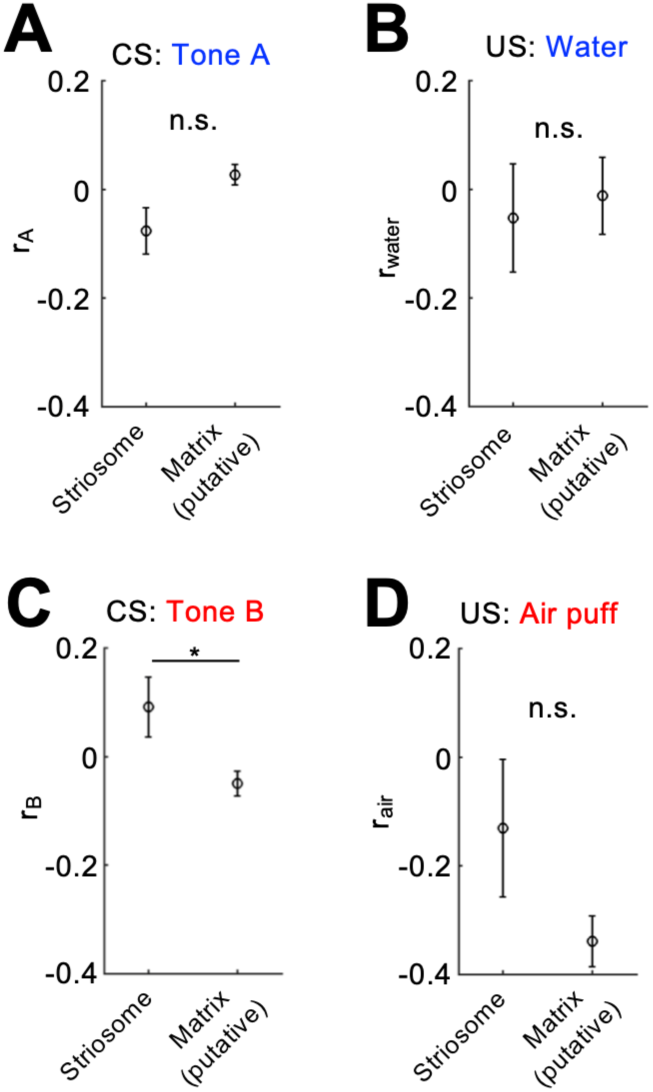
Different learning effects between striosomes and the putative matrix on DA release in response to Tone B. **A, B.** Correlation coefficients between the number of Tone A trials and DA release in response to Tone A (r_A_, **A**) and saccharin water (r_water_, **B**). The correlation coefficients between trial experience and DA release in response to both Tone A and saccharin water were not significantly different between striosomes and the putative matrix. Error bars indicate the SEM. n.s.: *p* ≥0.05, unpaired *t*-test. **C, D.** Correlation coefficients between the number of Tone B trials and DA release in response to Tone B (r_B_, **C**) and air puffs (r_air_, **D**). The correlation coefficient between trial experience and DA release in response to Tone B was significantly larger in striosomes than in the putative matrix, indicating a stronger learning effect in striosomes. *: *p* < 0.05.

## Discussion

We performed selective *in vivo* recordings of DA dynamics in striosomes during classical conditioning. To the best of our knowledge, this is the first report showing DA dynamics in striosomes in behaving animals. The major findings are as follows: (1) Air-puff-associated tones induced phasic DA release to striosomes; (2) In striosomes, DA release was induced by saccharin water intake and air puff stimuli; (3) In the putative matrix, saccharin water intake triggered augmentation of DA release, whereas air puff stimuli triggered suppression of DA release.

Although some studies have reported different DA dynamics between striosomes and the matrix in acute brain slices (Krebs et al., 1994; Krebs et al., 1991), it is unclear whether such a difference is observed in behaving animals. Because striosomes comprise only 15% of the striatum and are mosaically scattered, it is difficult to selectively record DA dynamics in striosomes using traditional neurochemical measurements, such as microdialysis or voltammetry. Therefore, it remains unclear how striosome-specific DA dynamics affect animal behavior. In this study, we successfully recorded DA release in striosomes during classical conditioning using fiber photometry in transgenic mice via selective DA sensor expression in striosomal neurons. This strategy can be extended to matrix-selective DA recordings using another transgenic mouse line expressing Cre in matrix neurons, such as calbindin-Cre mice (Evans et al., 2020). However, it was possible to make matrix-dominant DA measurements without transgenic mice because 85% of the entire striatum consists of matrix. Therefore, the DA dynamics recorded in mice expressing dLight1.1 in both striosomes and matrix were expected to be predominantly derived from the matrix.

A tracing study revealed the presence of two groups of dopaminergic neurons with dominant projections to the striosomes or matrix (Matsuda et al., 2009). Therefore, we speculated that if different firing patterns are observed in response to the CS and US, this would indicate that DA dynamics differ between striosomes and the matrix. In fact, dopaminergic neurons projecting to different striatal regions, such as the DMS and dorsolateral striatum, have different response properties to rewarding and aversive stimuli (Lerner et al., 2015). In this study, saccharin water was a reward for mice because water intake was restricted except when performing the behavioral task. However, air puff stimuli are generally used as aversive stimuli. The striosome- and matrix-projecting dopaminergic neurons possibly showed different spike activities in Tone B trials, resulting in unique DA release dynamics from their terminals. Some genetic studies have reported that striosomal neurons express the prodynorphin gene, a precursor of dynorphin (Cui et al., 2014; Gong et al., 2003). Dynorphin binds to kappa-opioid receptors expressed in the presynaptic membrane of dopaminergic neurons (James et al., 1982). Previous studies reported that dynorphin inhibits DA release to the striatum during chronic inflammatory pain (Narita et al., 2005; Petraschka et al., 2007; Suzuki et al., 1999). The unique DA dynamics in striosomes might be caused by dynorphin-induced activation of kappa-opioid receptors at presynaptic terminals of dopaminergic neurons. The prelimbic cortex (PL), a region of the medial prefrontal cortex, projects preferentially to striosomes in the DMS (Donoghue and Herkenham, 1986; Friedman et al., 2015; Gerfen, 1984). PL neurons are activated by air puff stimuli (Clarke et al., 2023; Somogyi et al., 2021), and their activation is important for fear conditioning (Corcoran and Quirk, 2007). Furthermore, optogenetic inhibition of axon terminals of PL neurons reduced sensitivity to aversive light exposure in a cost-benefit conflict situation (Friedman et al., 2015). Our striosome-selective DA recordings revealed that the air-puff-predictive stimulus induced DA release in striosomes. Cortico-striatal synapses show DA-dependent plasticity that is suitable for reinforcement learning (Reynolds et al., 2001). According to these observations, it is possible that the cortical input from the PL to striosomes was reinforced by DA, resulting in the air-puff-predictive activity in striosomal neurons (Yoshizawa et al., 2018). Reinforcement learning models of the cortico-basal ganglia postulate that striosomal neurons learn state values, whereas matrix neurons are involved in either action selection (actor) or action value learning (Barto, 1995; Doya, 2000, 2002). A subsequent question is whether neural activity and DA dynamics differ between striosomes and the matrix during decision making.

In the putative matrix, saccharin water intake and air puff presentation augmented and suppressed DA release to the DMS, respectively. This result is consistent with calcium dynamics in terminals of DMS-projecting dopaminergic neurons (Lerner et al., 2015). However, in striosomes, both USs induced DA release to striosomes in the DMS. Similar to DA dynamics in DMS striosomes, dopaminergic neurons projecting to the dorsolateral striatum respond to both reward and aversive stimuli and therefore encode stimulus saliency (Lerner et al., 2015). The unique DA dynamics in response to USs in striosomes might contribute to air-puff- and reward-responsive neural activities in striosomes (Yoshizawa et al., 2018).

## Acknowledgements

We thank Dr. Kenji Doya at Okinawa Institute of Science and Technology Graduate University for providing the experimental apparatus. We thank Edanz (https://jp.edanz.com/ac) for editing the English text of a draft of this manuscript.

## Author contributions

T.Y. and M.F. designed the study; T.Y. performed the study; T.Y. contributed unpublished reagents/analytic tools; T.Y. analyzed the data; T.Y. and M.F. wrote the manuscript.

## Conflict of Interest

The authors report no conflicts of interest.

## Funding sources

This work was supported by JSPS KAKENHI Grant Number JP22K15217 and the generous research support of Hokkaido University for young researchers.

## Notes

### Competing Interest Statement

The authors have declared no competing interest.

